# Connectivity in the Dorsal Visual Stream is Enhanced in Action Video Game Players

**DOI:** 10.1101/2024.09.26.615213

**Authors:** Kyle Cahill, Timothy Jordan, Mukesh Dhamala

## Abstract

Action video games foster competitive environments that demand rapid spatial navigation and decision-making. Action video gamers often exhibit faster response times and slightly improved accuracy in vision-based sensorimotor tasks. However, the underlying functional and structural changes in the two visual streams of the brain that may be contributing to these cognitive improvements have been unclear. Using functional and diffusion MRI data, this study investigated the differences in connectivity between gamers who play action video games and nongamers in the dorsal and ventral visual streams. We found that action video gamers have enhanced functional and structural connectivity, especially in the dorsal visual stream. Specifically, there is heightened functional connectivity—both undirected and directed—between the left Superior Occipital Gyrus and the left Superior Parietal Lobule during a moving-dots discrimination decision-making task. This increased connectivity correlates with response time in gamers. The structural connectivity, as quantified by diffusion fractional anisotropy and quantitative anisotropy measures of the axonal fiber pathways between the same regions was also enhanced for gamers compared to nongamers. These findings provide valuable insights into how action video gaming can induce targeted neuroplastic changes, enhancing structural and functional connectivity between specific brain regions in the visual processing pathways. These connectivity changes in the dorsal visual stream underpin the superior performance of action video gamers compared to non-gamers in tasks requiring rapid and accurate vision-based decision-making.

**GRAPHICAL ABSTRACT:** 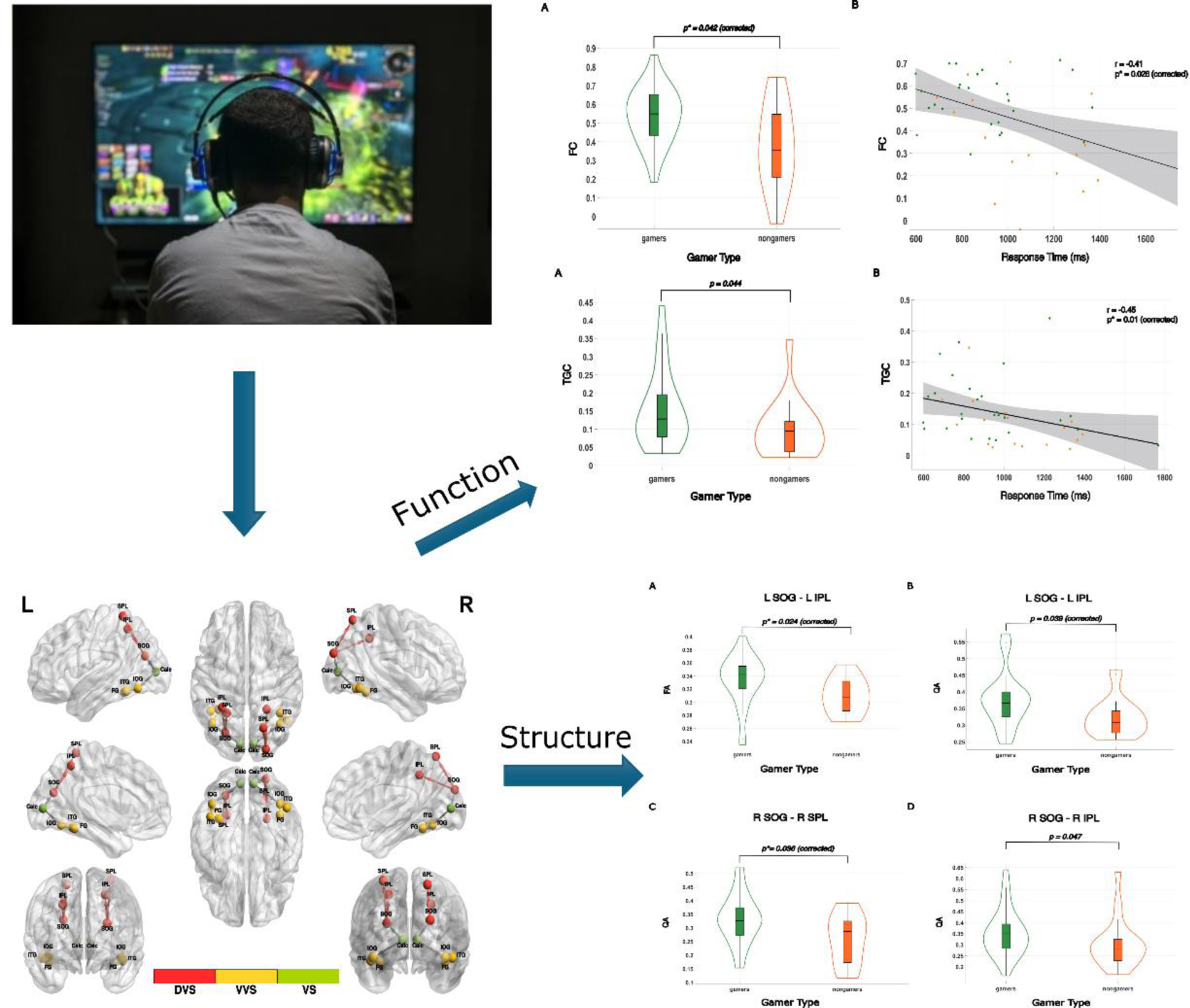

**Impact Statement:** Understanding the neural mechanisms underlying how observed cognitive alterations due to video game playing are achieved has potential implications for future cognitive training, rehabilitation, and education. The structural and functional MRI analysis in this study provides a quantitative basis by which the underlying neural connectivity changes due to video game playing in the visual streams may be assessed and linked to observed behavioral differences. These methods are extendible and provide insight into underlying neural network enhancements associated with improved cognitive performance due to video game playing. The findings of this study support enhanced functional and structural connectivity in the dorsal visual stream among gamers due to video game playing in the behavioral paradigm of vision-based sensorimotor decision-making. Functional connectivity measures that were considered were significantly correlated with the participants’ response times.

## INTRODUCTION

Action video games immerse the gamer in a fast-paced, challenging environment that heavily taxes vision-based decision-making. Gamers must rapidly process visual information and make swift yet accurate decisions, often refining their skills through many hours of dedicated play. While psychology and neuroscience literature have reported negative effects of video game playing such as the potential for video game addiction^1-3^ and increased aggression in those who play violent video games,^4,5^ there are also cognitive benefits of video game playing that are well-documented. Prolonged exposure to action video games is associated with improved cognitive performance, ranging from improvements in low-level perceptual decision-making to high-level cognitive flexibility.^6-14^ Action video gamers demonstrate better filtering of distracting information^15^, enhanced spatial attention^16^, improved accuracy in tracking multiple objects^17^, enhanced visual search^18,19^, reduced sensitivity to backward masking^20^, improved contrast sensitivity^21^, and mental rotation skills.^22^ In addition to these cognitive benefits, neuroimaging studies have demonstrated various structural and functional alterations in certain brain areas and associated brain networks among action video gamers. ^23-33^ However, only a few video game studies have conducted analysis on both structural and functional MRI data and even fewer have done so in healthy, non-addicted participants.^25,33^ Additionally, many previous studies, including those involving video game interventions, have not demonstrated how changes in neuroimaging metrics are reflected in cognitive changes due to a lack of cognitive tests in these studies to ascertain the relationship between cognitive function and neuroplastic changes. ^33^ This leaves results open-ended as it is inconclusive whether the neuroimaging changes found in one study can predict behavioral changes found in other studies and whether functional or structural measures are more effective as potential biomarkers.

The human brain is widely recognized as an intricate, complex biological system. A complete understanding of how it operates and adapts requires functional and structural analysis of its constituent subsystems.^34,35^ Establishing clear brain-behavior relationships, such as long-term action video game playing and the neuroplastic changes it induces, addresses a significant knowledge gap between behaviors and induced brain network-level changes. These neuroplastic changes may contribute to improvements in behavioral metrics ^36^ associated with cognitive processes, such as response times during vision-based sensorimotor decision-making. This highlights two important brain-behavior relationships: the connection between behaviors inducing neuroplastic changes, and the neuroplastic changes driving improvements in behavioral observables associated with specific cognitive processes due to engaging in those behaviors.

To help address this gap in the literature, this study investigates how long-term action video game play impacts both functional and structural connectivity in the two visual streams—the dorsal ‘where/how’ stream and the ventral ‘what’ stream—which are the networks most closely involved in visual processing as outlined by the well-established “two-streams” hypothesis.^37-45^ In a previous study using a low-level moving dots task designed to probe participants’ vision-based sensorimotor decision-making accuracy and response time, we found that action video gamers have faster response times (∼190ms) without compromising accuracy.^13,46^ By examining the two visual streams, we aim to expand the current understanding of the impact of long-term action video game playing has on the neural mechanisms driving this improvement in vision-based sensorimotor decision response times by illuminating clear brain-behavior relationships between parts of the streams that show significant differences in connectivity metrics between gamer and nongamer cohorts. These brain-behavior relationships would potentially provide an effective biomarker that predicts a person’s vision-based sensorimotor decision-making response time.

In this study, we utilized both functional connectivity and structural connectivity measures. We used undirected functional connectivity (FC), also called a Pearson correlation, and a directed functional connectivity measure, time-domain Granger causality (TGC). As opposed to undirected FC analysis, TGC allowed for the determination of the directional influence of one region onto another. This method elucidates how visual information is processed through both visual streams and how this flow of information is altered for long-term game playing. From both measures, we were able to establish not only brain-behavior relationships with these functional measures but also determine if the directionality of information flows plays an important role in the observed behavioral benefits.

Our structural connectivity analysis used fractional anisotropy (FA), a standard diffusion tensor imaging measure based on diffusivity ^44^ and has known associations with “action per minute” in certain brain areas of participants who underwent training through playing the real-time strategy game, “Starcraft II” including the left Inferior Longitudinal Fasciculus.^29^ Additionally, we considered the quantitative anisotropy (QA) measure because QA-based tractography is known to outperform FA-based tractography and is derived from the Fourier transform relation between MR signal and diffusion displacement.^47^ Utilization of both of these measures allowed for a clear and intuitive characterization of the white matter organization and microstructures by leveraging diffusivity and diffusion displacement.

Given that action video games involve extensive spatial exploration, navigation, and rapid time coordination, we hypothesize that both functional and structural connectivity within the visual streams may undergo neuroplastic enhancements due to prolonged action video game playing. Furthermore, we posit that elevated brain connectivity metrics are likely related to the improved response times observed in gamers. This study addresses changes in neuroimaging metrics due to action video game playing and conclusively determines the effectiveness of each metric in predicting response times during vision-based, sensorimotor decision-making. By employing these neuroimaging methods, we aim to enhance the understanding of how long-term action video game playing affects connectivity in the visual streams and its relationship with response time, while also providing a framework for subsequent research in this field.

## 2. RESULTS

### 2.1 Functional Connectivity

#### 2.1.1 Elevated Functional Connectivity in Gamers and Correlation with Response Time

Pairwise Pearson correlation was utilized for undirected functional connectivity analysis. Statistical comparison between video gamers and nongamers was conducted using the Wilcoxon test. In comparison, gamers exhibited higher FC values, between the L SOG and the L SPL. This result survived Holm-Bonferroni multiple comparisons after accounting for four connections among the ROIs composing the dorsal stream and remained statistically significant at the 0.05 level, p* = 0.042. A violin plot was constructed to graphically illustrate the significant difference found in FC values between gamers and nongamers in Fig 1, A, FC values were plotted against RT, and a linear regression analysis, depicted in Fig 1, B, was employed to explore the existence of a brain-behavior link; FC was found to have a significant moderate level correlation with subjects’ response time (r = -0.41, p* = 0.026).

**Fig 1.**
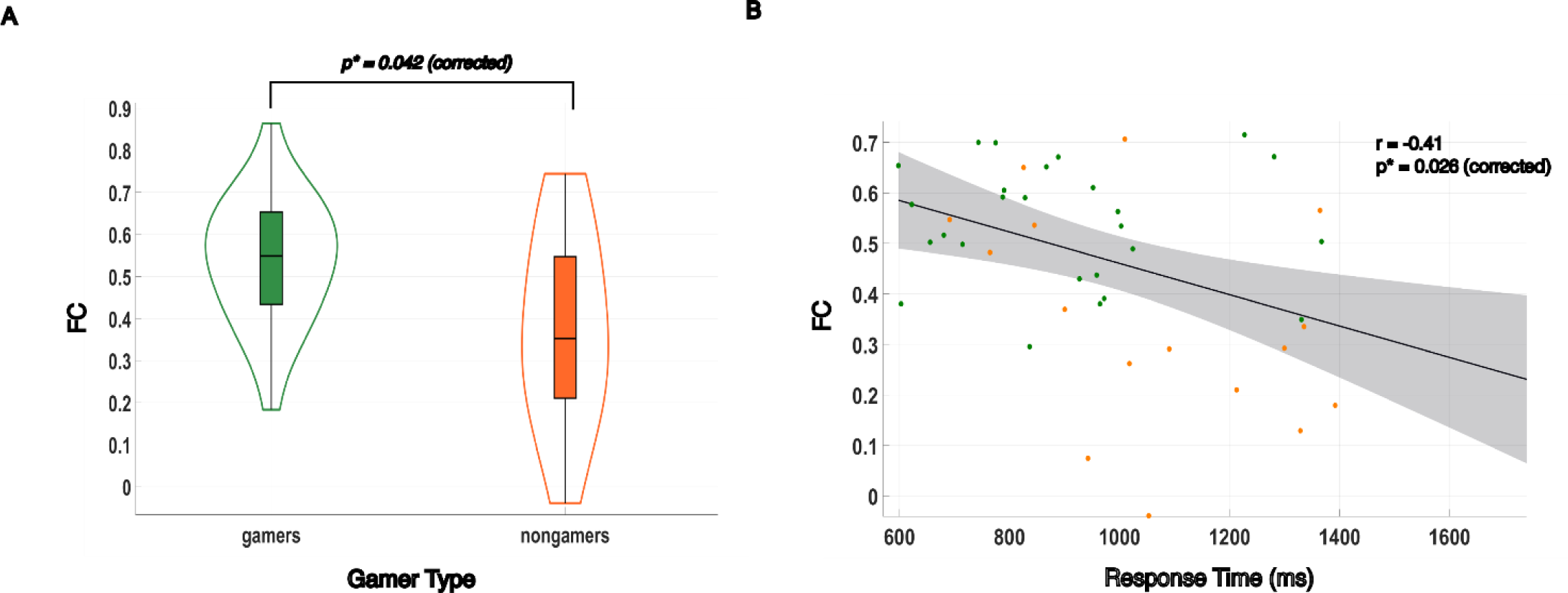
Undirected functional connectivity between the Left Superior Occipital Gyrus and Left Superior Parietal Lobule and brain-behavior correlation with response times. **A)** Gamers showed significantly higher FC values between the L SOG and the L SPL, p* = 0.042. **B)** The undirected functional connectivity measure was significantly correlated with response time, (r = -0.41, p* = 0.026). The gamers are indicated by the green dots while nongamers are indicated by the orange dots.

#### 2.1.2 Elevated Granger Causality in Gamers and Correlation with Response Time

The directed connectivity analysis utilized pairwise Granger causality to determine differences in directional influence along the visual streams. After testing a range of possible model orders through computation aimed at minimizing the total spectral difference between the Granger time series and the original, we found that the optimal model order for our data set was a model order of six. Time domain Granger causality (TGC) was computed for all pairwise links between subjects. Comparing gamers to nongamers, elevated TGC values were also observed between the L SOG and the L SPL with an uncorrected p-value, p = 0.044. This significant difference in TGC values is visually represented in Fig 2, A, through a violin plot. We further investigated the relationship between time-domain granger causality values and observed response times. TGC values were plotted against RT, and a linear regression analysis, depicted in Fig 2, B, was employed to explore the existence of a brain-behavior link; TGC was found to have a significant moderate level correlation with subjects’ response time (r = -0.45, p* = 0.01 (corrected)).

**Fig 2.**
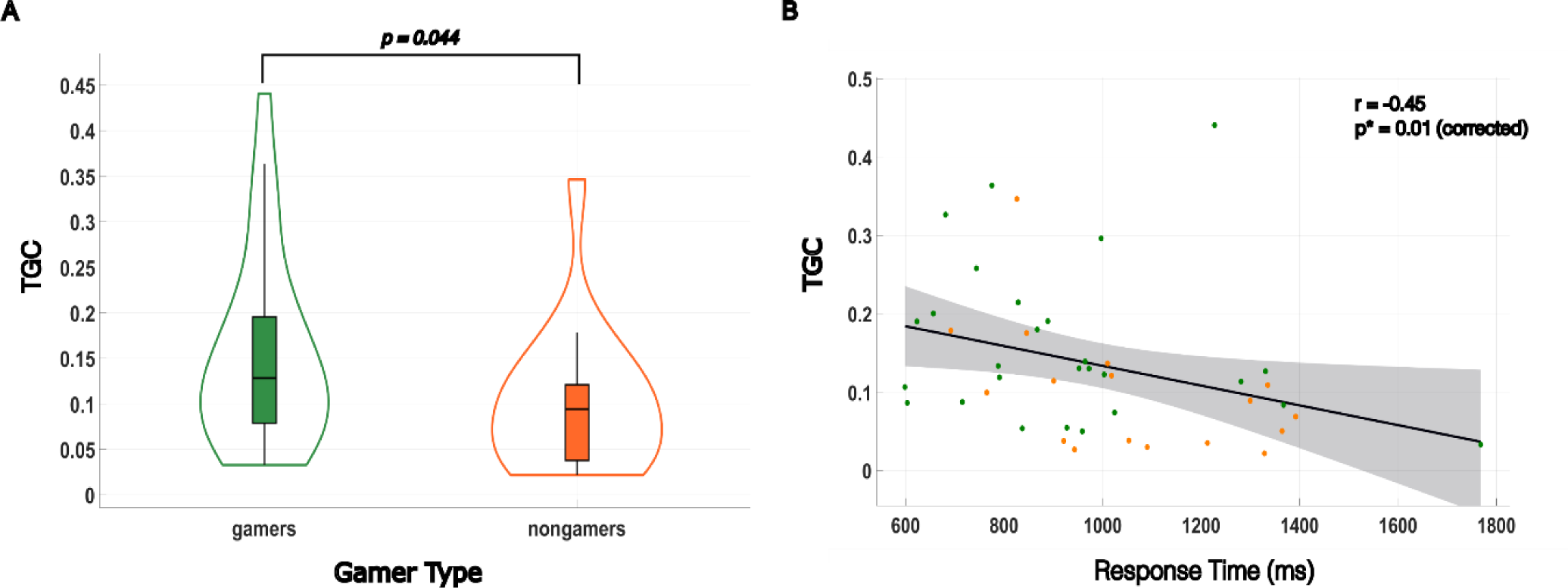
Directed functional connectivity, (TGC) between the Left Superior Occipital Gyrus (L SOG) and Left Superior Parietal Lobule (L SPL) and brain-behavior correlation with response times. **A)** Gamers showed significantly higher TGC values between the R SOG and the R SPL, uncorrected p=0.044. **B)** The directed functional connectivity measure TGC was significantly correlated with response time, (r = -0.45, p* = 0.01). The gamers are indicated by the green dots while nongamers are indicated by the orange dots.

### 2.2. Structural Connectivity

#### 2.2.1 Elevated Fractional Anisotropy and Quantitative Anisotropy in Gamers

The statistical comparison of structural connectivity measures, specifically fractional anisotropy and quantitative anisotropy between gamers and nongamers utilized the Wilcoxon test. In this analysis, gamers exhibited elevated FA values between the L SOG and the L IPL (p* = 0.024). Gamers also demonstrated elevated QA values between the same regions (L SOG and L IPL) with statistical significance (p* = 0.039). In addition to the observed elevation in FA and QA values within the left dorsal stream, notably between L SOG and L IPL, our investigation revealed heightened QA values in the right dorsal stream as well. Specifically, gamers exhibited increased QA R SOG and the R SPL (p* = 0.036), as well as between R SOG and the R IPL (p = 0.047). The increased QA between R SOG and R IPL did not survive multiple comparison corrections but remained significant at the individual level, warranting consideration for further study. To visually illustrate the significant differences in values between gamers and nongamers, violin plots were constructed (Fig 3). Structural connectivity measures FA and QA did not show a significant correlation with subjects’ response time.

**Fig 3.**
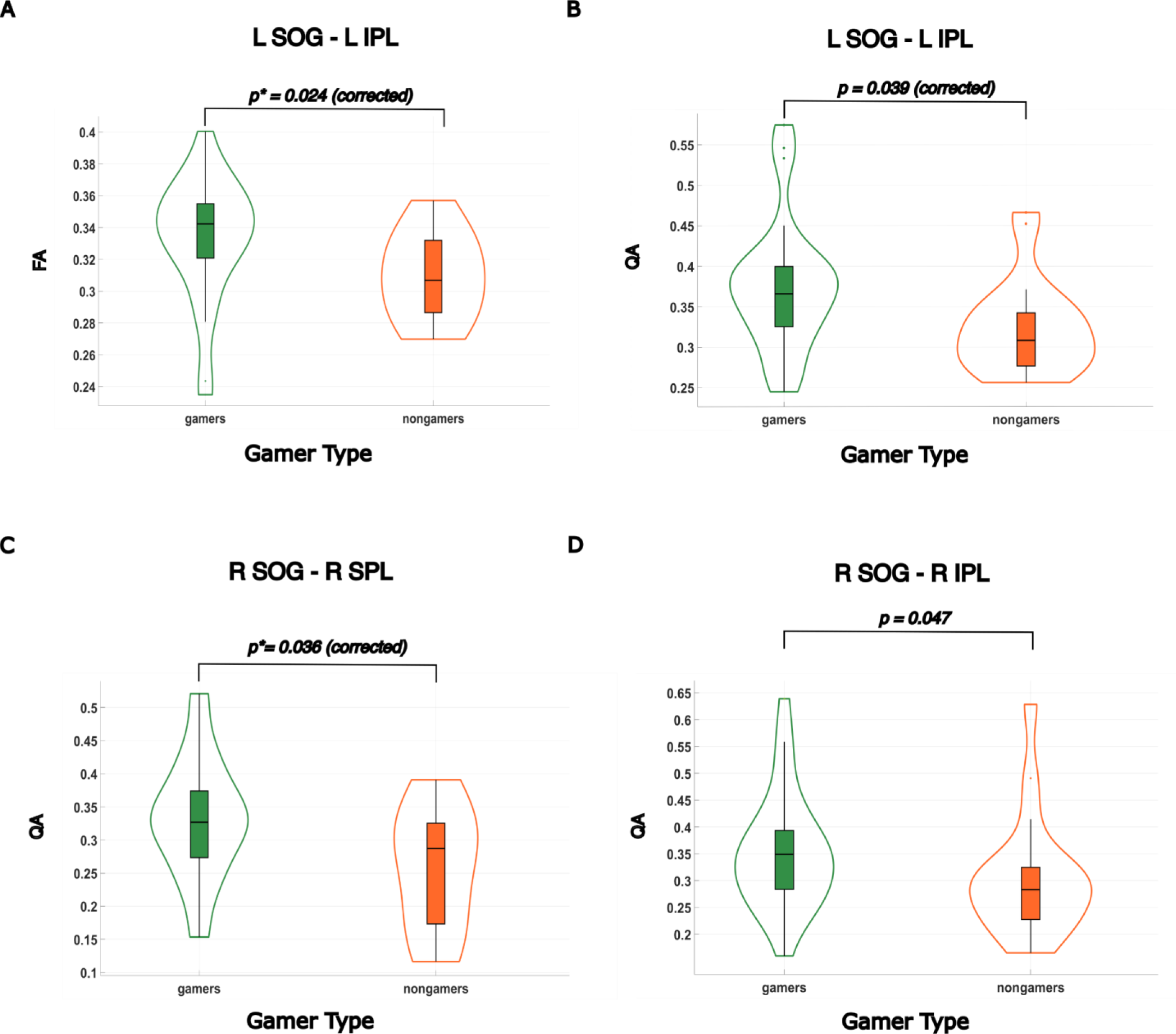
Structural connectivity in the dorsal stream is elevated in gamers. **A)** Gamers showed significantly higher FA values between the Left Superior Occipital Gyrus (L SOG) and the Left Inferior Parietal Lobule (L IPL), p*=0.024. **B)** Video gamers showed significantly higher QA values between the L SOG and the L IPL, p*=0.039. **C)** Video gamers showed significantly higher QA values between the Right Superior Occipital Gyrus (R SOG) and the Right Superior Parietal Lobule (R SPL), p*= 0.036. **D)** Video gamers showed significantly higher QA values between the R SOG and the Right Superior Parietal Lobule (R IPL), uncorrected p= 0.047.

## DISCUSSION

This study investigated functional and structural connectivity differences between gamers and nongamers in the dorsal and ventral visual streams. From our structural results, we observed that gamers exhibited increased white matter structural connectivity within the dorsal visual stream. Specifically, the connection in the left DVS between the left superior occipital gyrus and the left inferior parietal lobule showed elevated fractional anisotropy (FA) and quantitative anisotropy (QA) values, as well as elevated QA values in the right DVS. FA has known associations with axonal integrity and myelination, while QA is linked to axonal density. ^47-49^ These properties could indicate greater directional coherence and suggest better white matter fiber tract organization between brain areas. Although structural differences were observed, they did not correlate with response time.

We also observed higher functional connectivity values in the left DVS for gamers compared to nongamers. We found that undirected functional connectivity values and time-domain Granger causality values were higher between the left superior occipital gyrus and the left superior parietal lobule. Both measures of connectivity showed moderate correlations with response times, reflecting that they may play a strong role in influencing gamer performance specifically in response times. Taken together, these results show that extensive video game playing does alter both structural and functional connectivity in gamers which supports previous findings. However, our study shows that the functional connectivity measures FC and TGC serve as better gauges of participants’ response times compared to the structural connectivity measures FA and QA during vision-based sensorimotor decision-making. While the elevated TGC values didn’t survive corrections for multiple comparisons, this should be further explored in future studies due to its correlation with performance.

This study has some limitations, including an imbalance in the sex distribution between cohorts^50^, and the focus on healthy, age-matched young adults ^51-54^ which could potentially influence the generalizability of our findings. Future research should address these limitations with a more diverse participant group. While our study’s sample size was sufficient for statistical comparison, larger participant pools in future studies would increase overall statistical power and thus improve generalizability. FA has limitations in that this measure is known to suffer from partial volume effects due to crossing fibers and free water in the brain.^55^ Consideration of these known weaknesses in FA-based tractography informed our decision to implement QA-based tractography to conduct the structural connectivity analysis since QA-based tractography is less susceptible to these confounding effects while still recording DSI Studio’s estimation of FA between ROIs.^55^ Lastly, this study combined all gamers into one group instead of separating them into their respective genres. Future research should continue to explore the impact of specific games, game genres, and subgenres on specific cognitive processes and their effects on brain connectivity to better understand the mechanisms driving cognitive improvements similar to some previous studies. ^56,57^

A crucial aspect of advancing our understanding of neuroplasticity due to video game playing involves continued research into both the rate at which structural and functional connectivity changes occur among gamers and the identification of key factors that may influence this rate. Insight into the timeline for these connectivity changes in response to gaming experiences and the relevant factors affecting this timeline could reveal critical thresholds and strategies for optimizing skill acquisition and windows for skill transfer. ^58^ ^59^ The analysis presented in this study can be extended to investigate possible connectivity differences in interstream interactions between gamers and nongamers. Previous work suggests that interactions between the dorsal and ventral streams are necessary for skilled grasp ^60^ and complex object manipulation,^61^ which gamers engage in frequently while playing an action video game. The analysis presented in this study can be extended to investigate possible connectivity differences in interstream interactions between gamers and nongamers. Finally, we plan to expand the functional and structural connectivity methods implemented in this study to the whole brain and expand upon our understanding of brain-wide neuroplastic changes due to long-term exposure to action video game playing.

Our findings offer robust evidence of enhanced structural and functional connectivity within the dorsal visual stream, highlighting the influence of video game experience on neuroplasticity. Prior research has established the dorsal stream’s involvement in tracking object location and trajectories in space,^38^ which is vital information in action video games. The increased functional connectivity measures FC and TGC between the Left Superior Occipital Gyrus (L SOG) and the Left Superior Parietal Lobule (L SPL), correlate with improved sensorimotor decision-making response times, indicating that this connection within the ‘where/how’ pathway is linked to better functional visual-information processing thereby likely driving improvements in response times during vision-based sensorimotor decision making.

## Conclusion

In summary, this study provides compelling evidence that prolonged action video game exposure enhances both structural and functional connectivity in the dorsal visual stream. Specifically, the enhanced functional connectivity measures between the left superior occipital gyrus and the left superior parietal lobule are associated with improved response times during sensorimotor decision-making. These results suggest that video game experience can drive significant neuroplastic changes linked to cognitive improvements. Consequently, these findings could significantly impact cognitive training, rehabilitation, and educational strategies, offering new insights into the potential for video games to promote neuroplasticity and cognitive enhancement. This study makes a significant contribution by examining the impact of long-term action video game exposure on measurable neuroplastic changes in visual stream connectivity.

The dual-metric framework used in this study provides a comprehensive and intuitive method for quantifying the extent to which behavioral experiences influence neural connectivity and how these changes relate to cognitive performance. This methodological advancement establishes a solid foundation for future research into the effects of various video game genres on cognitive processes, highlighting the potential of video game studies as a valuable tool for investigating behaviorally induced neuroplasticity. ^62^ Furthermore, the methods employed in this MRI analysis present a rigorous approach for assessing neural connectivity changes due to video game playing and linking these changes to observable behavioral improvements.

## 4. Materials & Methods

### 4.1 Subject Data

47 participants (28 gamers (4 female) and 19 nongamers (12 female)) were recruited to participate. Groups were age matched (gamers = 20.6 ± 2.4 years, nongamers = 19.9 ± 2.6 years). Participants who indicated playing 5 hours per week or more in one of four types of video game genres for the last two years were considered video game players i.e., gamers. The four types of action video game player genres we recruited, based on industry demographics, were First-Person Shooter (FPS), Real-Time Strategy (RTS), Multiplayer Online Battle Arena (MOBA), and Battle Royale (BR) players. Participants who were nongamers in this study averaged less than 30 min per week in any video game over the last two years. A modified left-right moving dots (MD) motion categorization task was used to probe for differences in reaction speed and accuracy between the cohorts.^13,27^ We excluded two subjects’ data (one gamer and one nongamer) from the brain-behavior regressions due to incomplete response time data, one gamer from the structural connectivity analysis due to incomplete tractography data, and one additional nongamer from the functional connectivity analysis due to incomplete fMRI data.

### 4.2 MRI Data

Whole-brain structural and functional MR imaging was conducted on a 3 T Siemens Magnetom <u>Prisma</u> MRI scanner at the joint Georgia State University and Georgia Institute of Technology Center for Advanced <u>Brain Imaging</u>, Atlanta, Georgia. First, high-resolution anatomical images were acquired for voxel-based morphometry and anatomical reference using a T1-MEMPRAGE scan sequence (TR = 2530 ms, TE1-4: 1.69–7.27 ms, TI = 1260 ms, Flip Angle = 7 deg, Voxel size 1 mm × 1 mm x 1 mm). Following, four functional runs used a T2*-weighted gradient echo-planar imaging sequence (TR = 535 ms, TE = 30 ms, Flip Angle = 46°, Field of View = 240 mm, Voxel Size = 3.8 mm × 3.8 mm x 4 mm, Number of slices = 32 collected in an interleaved order, Slice thickness = 4 mm) and acquired 3440 brain images while participants performed the behavioral tasks. ^13,27^

### 4.3 Regions of Interest

In our investigation, we examined the structural and functional organization of the dorsal and ventral visual streams using fourteen regions of interest (ROIs) defined in a prior study from the Neurosynth functional meta-analysis platform (https://neurosynth.org/). The ROIs were derived based on relevant search terms, including “primary visual,” “ventral visual,” “visual stream,” and “dorsal visual.” ^63^ The identified ROIs encompassed two regions situated within the primary visual cortex (V1), specifically the bilateral Calcarine (Calc) areas. VS’ (visual stream) denotes the primary connections from the Calcarine region to the bilateral superior and inferior occipital gyri. Additionally, we identified four ROIs within the ventral visual stream (VVS), which included the bilateral fusiform gyrus (FG), and inferior temporal gyrus (ITG). Moreover, four ROIs were identified in the dorsal visual stream (DVS), encompassing the bilateral inferior parietal lobule (IPL), and superior parietal lobule (SPL). The nomenclature used in this classification was based on the Eickhoff-Zilles macro labels from N27 and was implemented in AFNI. ^63^We constructed a BrainNet Viewer^64^ representation of the general organization, subsystems, and the 14 ROIs along with the 12 connections composing the visual streams (Fig. 4).

**Fig. 4:**
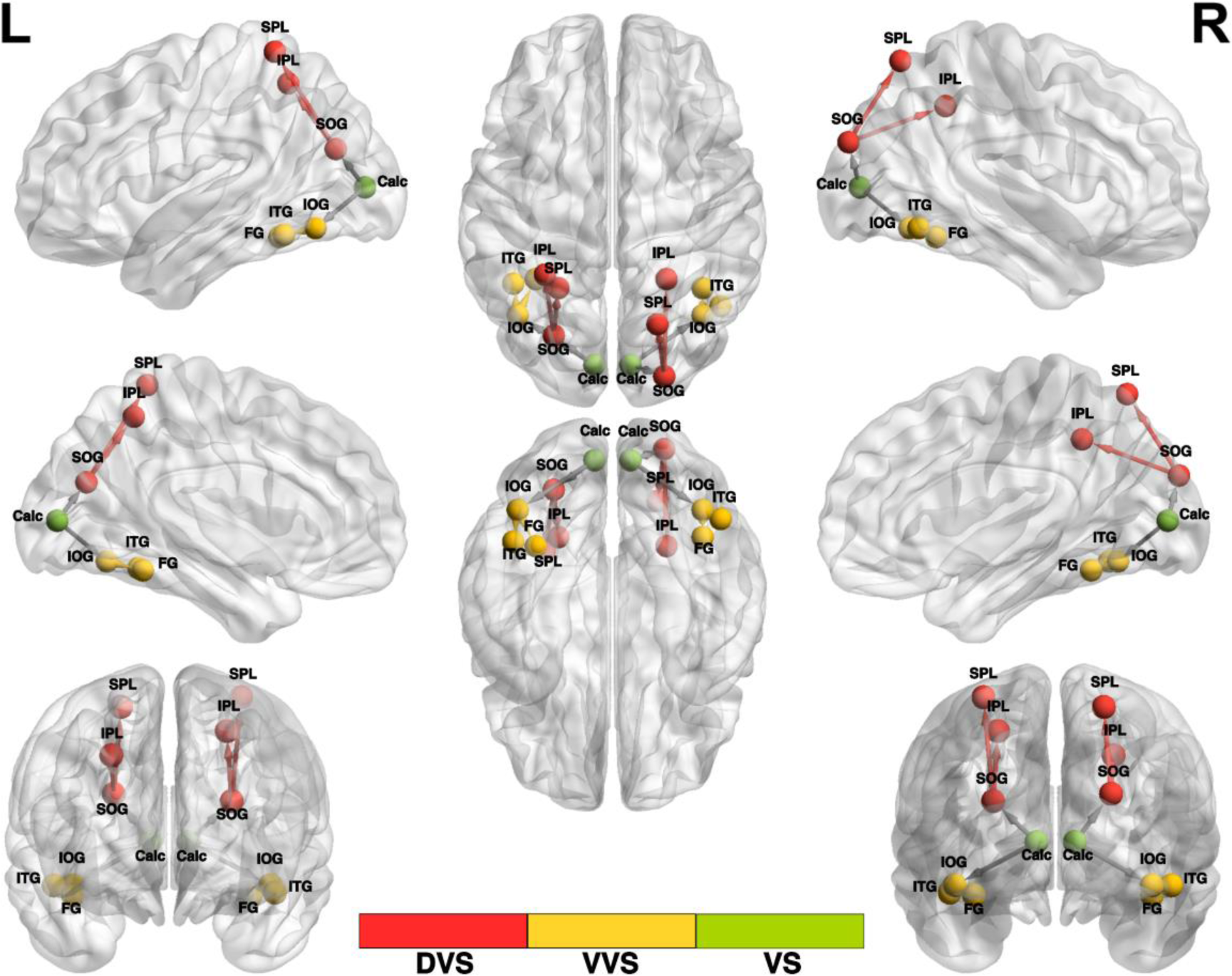
Meta-analysis derived ROIs for the dorsal and ventral visual streams. BrainNet Viewer representation of the locations of the 14 spherical ROIs and 12 connections considered that constitute the subsystems of the two visual streams. (**Left to Right):** The dorsal visual stream (**DVS**) extends from SOG to IPL and from SOG to SPL, shown in red; the ventral visual stream (**VVS**) extends from IOG to FG and from IOG to ITG, shown in yellow; and the visual stream (**VS**) denotes the primary connections from the Calcarine region namely Calcarine to IOG and Calcarine to SOG.

The construction of the ROIs for the structural tractography was carried out with a 12 mm radius to ensure the analysis accounted for anatomical variability and adequately encompassed the white matter tracts connecting the visual streams thus mitigating the risk of missing important connections. We employed the MNI coordinate system and constructed the ROIs using the FSLeyes visualization tool within the FSL (FMRIB Software Library) environment. The functional connectivity analysis utilized 6 mm radius ROIs using the same MNI coordinates.

For multiple comparison corrections in this analysis, we employed the Holm-Bonferroni method. This method was selected for its ability to enhance statistical power and sensitivity to individual significant comparisons while effectively controlling for Type I errors.^65,66^ We applied a significance threshold of p < 0.05, Holm-Bonferroni corrected indicated by p*, within each main section (DVS, VVS, VS) to ensure statistical significance while simultaneously controlling for the family-wise error rate in the dorsal and ventral streams, providing a robust framework for identifying meaningful differences in connectivity metrics.

### 4.4 Functional Connectivity Protocols and Analysis

Functional connectivity in neuroimaging measures temporal correlations of the BOLD signal time series in spatially distant brain regions.^67-69^ Thus, two regions are considered functionally connected if there is a statistical relationship between the measures of recorded BOLD activity.^70^ Region- and task condition-specific fMRI time series segments were normalized, voxel averaged, and corrected for linear trends. To assess each participant’s region-to-region undirected functional connectivity (FC) i.e., pairwise Pearson correlation coefficients were computed using 6 mm spherical ROIs.

The appropriate model order for the Granger causality (GC) analysis was determined by minimizing the spectral difference between the Granger time series and the original time series. After evaluating model orders ranging from two to twenty, a model order of six was selected as it best minimized the spectral difference in our dataset. GC matrices were then computed using this optimal model order.

The GC from region 2 to region 1 in the frequency domain is defined as follows:

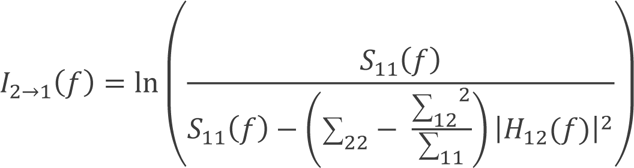

The frequency band-specific or time-domain equivalent GC is as follows:

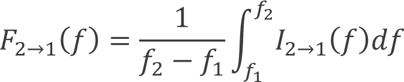

In the bivariate autoregressive model ^71-73^, the noise covariance and transfer function matrices are denoted by “∑” and “H” respectively. The evaluation of the frequency band-specific or time-domain equivalent Granger causality (GC) was accessed in the range between *f*_1_= 0.05 Hz to *f*_2_= 0.9 Hz, with a sampling rate of 1.87 Hz (*TR*^−1^)

To investigate the relationship between brain connectivity and behavior, undirected FC values and directed time-domain Granger causality (TGC) values were obtained for each participant per connection. Spearman’s rank correlation coefficient was used to assess the correlation between FC and reaction time (RT) as well as between TGC and RT.

### 4.5 Tractography Protocols & Structural Connectivity Analysis

*DSI Studio* is a non-commercial software that was utilized in this study for diffusion MR image analysis and provided functions including deterministic fiber tracking and 3D visualization.^55^ In this study, we used a multi-shell diffusion scheme with b-values of 300, 650, 1000, and 2000 s/mm². The acquisition parameters consisted of an in-plane resolution of 2 mm and a slice thickness of 2 mm. The accuracy of b-table orientation was examined by comparing fiber orientations with those of a population-averaged template ^74^. The diffusion data were reconstructed in the MNI space using q-space diffeomorphic reconstruction ^75^ to obtain the spin distribution function ^55,76^ A diffusion sampling length ratio of 1.25 was used. The resulting diffeomorphic reconstruction output had an isotopic resolution of 2 mm. Tensor metrics were calculated and a deterministic fiber tracking algorithm was used to track the fiber pathways.

Seeds were randomly placed throughout the ROIs until reaching a cutoff at 50,000,000 seeds. Additionally, two pairwise spherical ROIs were also defined as ending regions. In the case between the Left Superior Occipital Gyrus (L SOG) and the Left Inferior Parietal Lobule (L IPL), for example, the ending regions were placed at (52,74,37) and (51,64,51). An angular threshold of 60 degrees was set as the maximum allowed angular deviation between steps. The step size was randomly selected from 0.5 voxels to 1.5 voxels. Tracks with lengths shorter than 10 mm or longer than 100 mm were excluded from further analysis.

The process continued until mapping each subsystem of the dorsal and ventral visual streams (DVS, VVS, VS), with an exhaustive exploration of all pairwise links in each section. Adjustments to the parameters for maximum length and angular threshold were made based on the connection being mapped, as detailed in Table 2. For tracking eligibility, a quantitative anisotropy threshold of 0.01 was universally applied in all connections, except for the connection between R SOG and R IPL. In this case, a lower quantitative anisotropy threshold of 0.005 was necessary to prevent the inadvertent exclusion of subjects with valid fibers from the analysis. Consequently, voxels with a qualitative anisotropy value exceeding the specified threshold were deemed anisotropic and deemed suitable for inclusion in the fiber tracking process. The fiber pathways between the L SOG and L IPL are shown in a representative subject in Fig. 5 where the axis is color-coded to distinguish the orientation of the fibers. The X-axis is coded for red from right-left, the Y-axis is coded for green from anterior-posterior, and the Z-axis is coded for blue from superior-inferior.

**Fig. 5:**
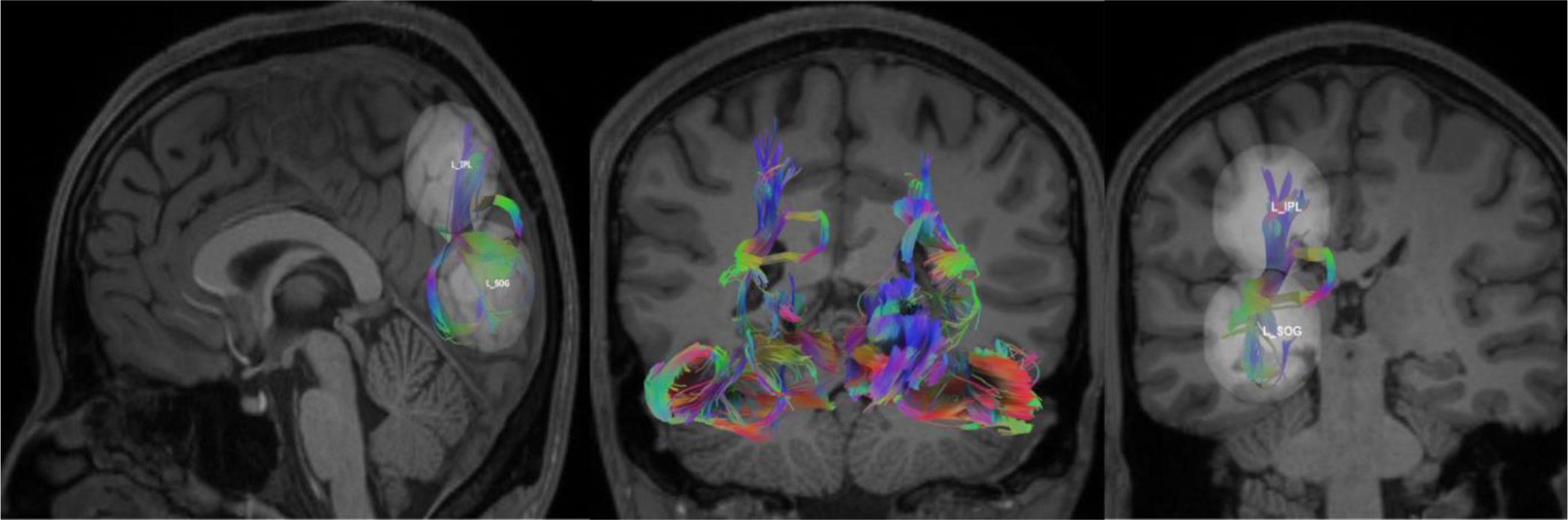
L SOG – L IPL fiber tracts overlayed on a T1 image in a representative participant. The fibers are colored-coded to represent their orientation, where “red” indicates fibers along the X-axis (i.e., left-right), “green” indicates fibers along the Y-axis (i.e., anterior-posterior), and “blue” indicates fibers along the Z-axis (i.e., inferior-superior). (**Left to Right): Sagittal,** reconstruction of white matter fiber tracts modelling pathways between L SOG and L IPL; **Coronal,** reconstruction of white matter tracts modelling the pathways that constitute the entire dorsal and ventral streams; **Coronal**, reconstruction of white matter tracts modelling the pathways between the L SOG and L IPL.

**Table 1:**
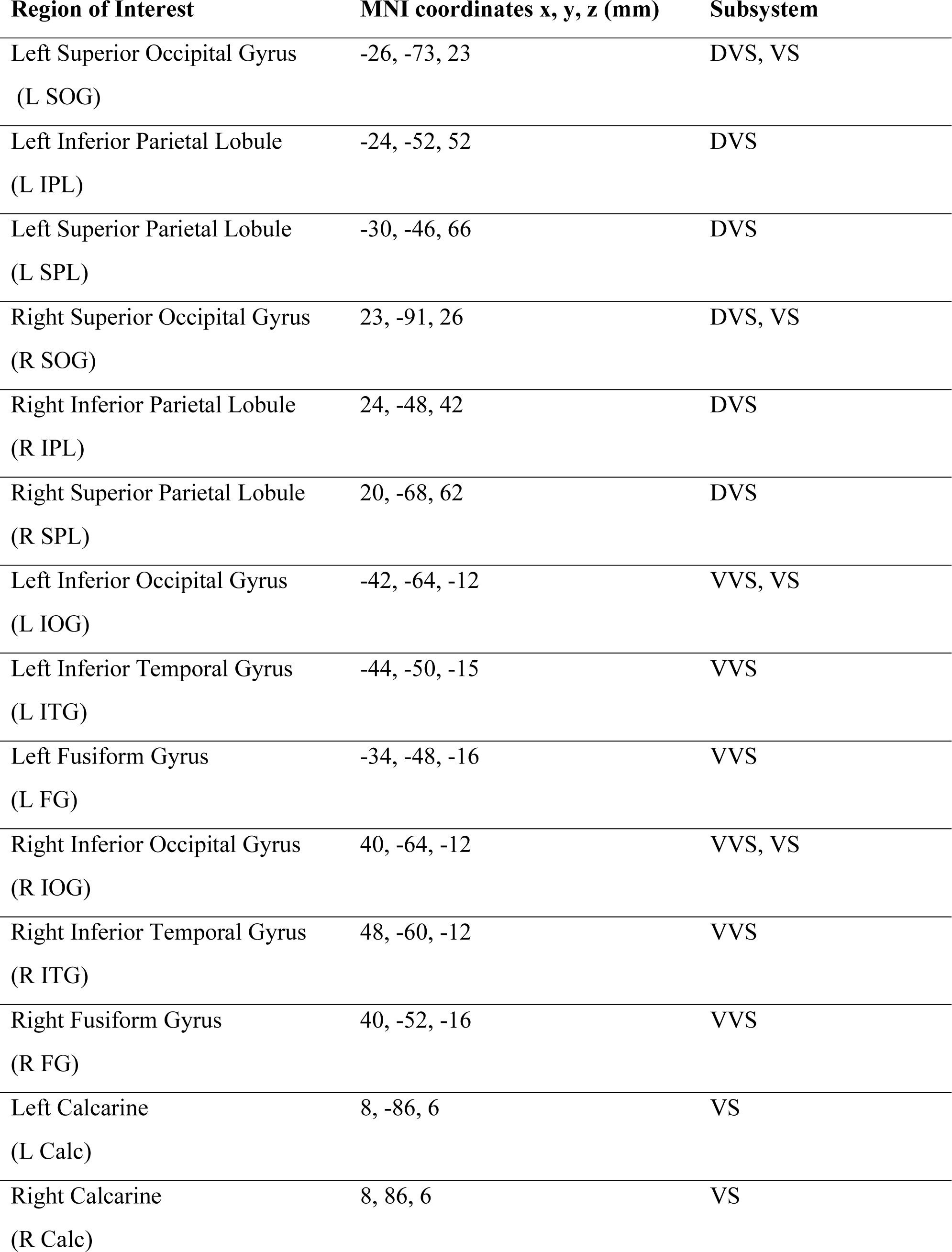
Regions of interest categorized by subsystem.

| **Region of Interest** | **MNI coordinates x, y, z (mm)** | **Subsystem** |
| --- | --- | --- |
| Left Superior Occipital Gyrus<br>(L SOG) | -26, -73, 23 | DVS, VS |
| Left Inferior Parietal Lobule<br>(L IPL) | -24, -52, 52 | DVS |
| Left Superior Parietal Lobule<br>(L SPL) | -30, -46, 66 | DVS |
| Right Superior Occipital Gyrus<br>(R SOG) | 23, -91, 26 | DVS, VS |
| Right Inferior Parietal Lobule<br>(R IPL) | 24, -48, 42 | DVS |
| Right Superior Parietal Lobule<br>(R SPL) | 20, -68, 62 | DVS |
| Left Inferior Occipital Gyrus<br>(L IOG) | -42, -64, -12 | VVS, VS |
| Left Inferior Temporal Gyrus<br>(L ITG) | -44, -50, -15 | VVS |
| Left Fusiform Gyrus<br>(L FG) | -34, -48, -16 | VVS |
| Right Inferior Occipital Gyrus<br>(R IOG) | 40, -64, -12 | VVS, VS |
| Right Inferior Temporal Gyrus<br>(R ITG) | 48, -60, -12 | VVS |
| Right Fusiform Gyrus<br>(R FG) | 40, -52, -16 | VVS |
| Left Calcarine<br>(L Calc) | 8, -86, 6 | VS |
| Right Calcarine<br>(R Calc) | 8, 86, 6 | VS |

**Table 2:** Tracking parameters for connections within each subsystem.

| Subsystem | Connection | Min length (mm) | Max length (mm) | Angular Threshold (deg) |
| --- | --- | --- | --- | --- |
| **DVS** | L SOG L IPL | 10 | 100 | 60 |
|  | L SOG L SPL | 10 | 300 | 70 |
|  | R SOG R IPL | 10 | 100 | 75 |
|  | R SOG R SPL | 10 | 150 | 70 |
| **VVS** | L IOG L ITG | 10 | 20 | 50 |
|  | L IOG L FG | 10 | 35 | 50 |
|  | R IOG R ITG | 10 | 20 | 50 |
|  | R IOG R FG | 10 | 35 | 50 |
| **VS** | L Calc L IOG | 10 | 70 | 65 |
|  | L Calc L SOG | 5 | 20 | 50 |
|  | R Calc R IOG | 10 | 80 | 65 |
|  | R Calc R SOG | 10 | 30 | 50 |

Structural connectivity assesses the anatomical organization of the brain using white matter fiber tracts.^77^ Although relatively stable on shorter time scales (seconds to minutes), it can exhibit plastic experience-dependent changes at longer periods (hours to days). ^78^ Fractional Anisotropy (FA), based on diffusivity, is calculated as the normalized fraction of the diffusion tensor’s magnitude.^79^ Fractional anisotropy (FA) is a useful measure and is often a standard in structural connectivity analysis.^79,80^ QA is known to be less susceptible to partial volume effects due to crossing fibers and free water in the brain than FA, resulting in a better resolution and improved tractography.^75,81^

Quantitative Anisotropy (QA) is a model-free measure derived from the Fourier transform relation between MR signals and diffusion displacement and is nonparametrically calculated from peak orientations on a spin distribution function.^55^ QA is known to be less susceptible to partial volume effects due to crossing fibers and free water in the brain than FA, resulting in a better resolution and improved tractography.^75,81^ Both fractional anisotropy (FA) and quantitative anisotropy (QA) were utilized as primary measures of enhanced structural connectivity and were extracted for each pair of ROIs composing the two visual streams and we investigated the brain-behavior relation of these two structural connectivity metrics with the participants’ response times.

#### Statistical analyses

Wilcoxon rank-sum tests on the connectivity measures and Pearson correlation analysis between the connectivity measures and response times were performed in MATLAB (The MathWorks). All tests are corrected for multiple comparisons using the Holm-Bonferroni procedure except for the sign-tests on Pattern coherence levels which were corrected using the Benjamini–Hochberg procedure.

## DATA AND CODE AVAILABILITY

All data that support the findings of the study as well as the custom analysis scripts can be found in OSF (*link will be public after publication*).

## ACKNOWLEDGEMENTS

This work was funded by two internal grant awards of Brains and Behavior Program & the Center for Advanced Brain Imaging to M.D.

## AUTHOR CONTRIBUTIONS

**Kyle Cahill:** Conceptualization, Methodology, Software, Formal analysis, Writing - original draft, review & editing.

**Mukesh Dhamala:** Conceptualization, Methodology, Software, Supervision, Funding acquisition, Writing - original draft, review & editing.

**Timothy Jordan:** Conceptualization, Methodology, review & editing

## Abbreviation List

AFNI: Analysis of Functional NeuroImages
BOLD: Blood Oxygen Level Dependent
BR: Battle Royale
Calc: Calcarine
DSI: Diffusion Spectral Imaging
DTI: Diffusion Tensor Imaging
FA: Fractional Anisotropy
FC: Functional Connectivity
FG: Fusiform Gyrus
fMRI: Functional Magnetic Resonance Imaging
FPS: First-Person Shooter
FSL: FMRIB Software Library
GC: Granger Causality
IOG: Inferior Occipital Gyrus
IPL: Inferior Parietal Lobule
ITG: Inferior Temporal Gyrus
L: Left
MD: Moving Dots
MNI: Montreal Neurological Institute
MOBA: Multiplayer Online Battle Arena
QA: Quantitative Anisotropy
R: Right
ROI: Region of Interest
RT: Response Time
RTS: Real-Time Strategy
SOG: Superior Occipital Gyrus
TI: Inversion Time
TR: Repetition Time

## REFERENCES

1 Mohammad, S., Jan, R. A. & Alsaedi, S. L. Symptoms, Mechanisms, and Treatments of Video Game Addiction. Cureus 15, e36957 (2023). 10.7759/cureus.36957

2 Gros, L., Debue, N., Lete, J. & van de Leemput, C. Video Game Addiction and Emotional States: Possible Confusion Between Pleasure and Happiness? Front Psychol 10, 2894 (2019). 10.3389/fpsyg.2019.02894

3 Weinstein, A., Livny, A. & Weizman, A. New developments in brain research of internet and gaming disorder. Neuroscience & Biobehavioral Reviews 75, 314–330 (2017). 10.1016/j.neubiorev.2017.01.040

4 Greitemeyer, T. The contagious impact of playing violent video games on aggression: Longitudinal evidence. Aggress Behav 45, 635–642 (2019). 10.1002/ab.21857

5 Yao, M., Zhou, Y., Li, J. & Gao, X. Violent video games exposure and aggression: The role of moral disengagement, anger, hostility, and disinhibition. Aggress Behav 45, 662–670 (2019). 10.1002/ab.21860

6 Powers, K. L., Brooks, P. J., Aldrich, N. J., Palladino, M. A. & Alfieri, L. Effects of video-game play on information processing: a meta-analytic investigation. Psychon Bull Rev 20, 1055–1079 (2013). 10.3758/s13423-013-0418-z

7 Dye, M. W., Green, C. S. & Bavelier, D. Increasing Speed of Processing With Action Video Games. Curr Dir Psychol Sci 18, 321–326 (2009). 10.1111/j.1467-8721.2009.01660.x

8 Dye, M. W., Green, C. S. & Bavelier, D. The development of attention skills in action video game players. Neuropsychologia 47, 1780–1789 (2009). 10.1016/j.neuropsychologia.2009.02.002

9 Green, C. S. & Bavelier, D. Action-video-game experience alters the spatial resolution of vision. Psychol Sci 18, 88–94 (2007). 10.1111/j.1467-9280.2007.01853.x

10 Green, C. S. & Bavelier, D. Action video game training for cognitive enhancement. Current Opinion in Behavioral Sciences 4, 103–108 (2015). 10.1016/j.cobeha.2015.04.012

11 Lynch, J., Aughwane, P. & Hammond, T. M. Video Games and Surgical Ability: A Literature Review. Journal of Surgical Education 67, 184–189 (2010). 10.1016/j.jsurg.2010.02.010

12 Basak, C., Boot, W. R., Voss, M. W. & Kramer, A. F. Can training in a real-time strategy video game attenuate cognitive decline in older adults? Psychol Aging 23, 765–777 (2008). 10.1037/a0013494

13 Jordan, T. & Dhamala, M. Video game players have improved decision-making abilities and enhanced brain activities. Neuroimage: Reports 2, 100112 (2022). 10.1016/j.ynirp.2022.100112

14 Howard, J., Bowden, V. K. & Visser, T. Do action video games make safer drivers? The effects of video game experience on simulated driving performance. Transportation Research Part F: Traffic Psychology and Behaviour 97, 170–180 (2023). 10.1016/j.trf.2023.07.006

15 Bavelier, D., Achtman, R. L., Mani, M. & Föcker, J. Neural bases of selective attention in action video game players. Vision Research 61, 132–143 (2012). 10.1016/j.visres.2011.08.007

16 Green, C. S. & Bavelier, D. Effect of action video games on the spatial distribution of visuospatial attention. J Exp Psychol Hum Percept Perform 32, 1465–1478 (2006). 10.1037/0096-1523.32.6.1465

17 Green, C. S. & Bavelier, D. Action video game modifies visual selective attention. Nature 423, 534–537 (2003). 10.1038/nature01647

18 Chisholm, J. D. & Kingstone, A. Action video game players’ visual search advantage extends to biologically relevant stimuli. Acta Psychologica 159, 93–99 (2015). 10.1016/j.actpsy.2015.06.001

19 Wu, S. & Spence, I. Playing shooter and driving videogames improves top-down guidance in visual search. Atten Percept Psychophys 75, 673–686 (2013). 10.3758/s13414-013-0440-2

20 Li, R., Polat, U., Scalzo, F. & Bavelier, D. Reducing backward masking through action game training. Journal of Vision 10, 33–33 (2010). 10.1167/10.14.33

21 Li, R., Polat, U., Makous, W. & Bavelier, D. Enhancing the contrast sensitivity function through action video game training. Nature Neuroscience 12, 549–551 (2009). 10.1038/nn.2296

22 Anguera JA, B. J., Rintoul JL, Al-Hashimi O, Faraji F, Janowich J, Kong E, Larraburo Y, Rolle C, Johnston E, Gazzaley A. Video game training enhances cognitive control in older adults. Nature. 2013 Sep 5;501(7465):97-101. doi: 10.1038/nature12486. PMID: 24005416; PMCID: PMC3983066.

23 Basak, C., Voss, M. W., Erickson, K. I., Boot, W. R. & Kramer, A. F. Regional differences in brain volume predict the acquisition of skill in a complex real-time strategy videogame. Brain Cogn 76, 407–414 (2011). 10.1016/j.bandc.2011.03.017

24 Kowalczyk, N. et al. Real-time strategy video game experience and structural connectivity – A diffusion tensor imaging study. Human Brain Mapping 39, 3742–3758 (2018). 10.1002/hbm.24208

25 Palaus, M., Marron, E. M., Viejo-Sobera, R. & Redolar-Ripoll, D. Neural Basis of Video Gaming: A Systematic Review. Frontiers in Human Neuroscience 11 (2017). 10.3389/fnhum.2017.00248

26 Nikolaidis, A., Voss, M., Lee, H., Vo, L. & Kramer, A. Parietal plasticity after training with a complex video game is associated with individual differences in improvements in an untrained working memory task. Frontiers in Human Neuroscience 8 (2014). 10.3389/fnhum.2014.00169

27 Jordan, T. & Dhamala, M. Enhanced Dorsal Attention Network to Salience Network Interaction in Video Gamers During Sensorimotor Decision-Making Tasks. Brain Connect (2022). 10.1089/brain.2021.0193

28 Granek, J. A., Gorbet, D. J. & Sergio, L. E. Extensive video-game experience alters cortical networks for complex visuomotor transformations. Cortex 46, 1165–1177 (2010). 10.1016/j.cortex.2009.10.009

29 Lewandowska, P. et al. Association between real-time strategy video game learning outcomes and pre-training brain white matter structure: preliminary study. Scientific Reports 12, 20741 (2022). 10.1038/s41598-022-25099-0

30 Huang, H. & Cheng, C. The Benefits of Video Games on Brain Cognitive Function: A Systematic Review of Functional Magnetic Resonance Imaging Studies. Applied Sciences 12, 5561 (2022).

31 Kühn, S., Gallinat, J. & Mascherek, A. Effects of computer gaming on cognition, brain structure, and function: a critical reflection on existing literature Dialogues Clin Neurosci 21, 319–330 (2019). 10.31887/DCNS.2019.21.3/skuehn

32 Kühn, S. & Gallinat, J. Amount of lifetime video gaming is positively associated with entorhinal, hippocampal and occipital volume. Mol Psychiatry 19, 842–847 (2014). 10.1038/mp.2013.100

33 Brilliant, T. D., Nouchi, R. & Kawashima, R. Does Video Gaming Have Impacts on the Brain: Evidence from a Systematic Review. Brain Sci 9 (2019). 10.3390/brainsci9100251

34 Pessoa, L. Understanding brain networks and brain organization. Phys Life Rev 11, 400–435 (2014). 10.1016/j.plrev.2014.03.005

35 Lynn, C. W. & Bassett, D. S. The physics of brain network structure, function and control. Nature Reviews Physics 1, 318–332 (2019). 10.1038/s42254-019-0040-8

36 Schlinger, H. D. Behavior analysis and behavioral neuroscience. Front Hum Neurosci 9, 210 (2015). 10.3389/fnhum.2015.00210

37 McIntosh, R. S., T. (May 2009). “Two visual streams for perception and action: current trends”. Neuropsychologia. 47 (6): 1391–6. doi:10.1016/j.neuropsychologia.2009.02.009. PMID 19428404. S2CID 32937236.

38 Schenk, T. & McIntosh, R. D. Do we have independent visual streams for perception and action? Cognitive Neuroscience 1, 52–62 (2010). 10.1080/17588920903388950

39 Kravitz, D. J., Saleem, K. S., Baker, C. I. & Mishkin, M. A new neural framework for visuospatial processing. Nature Reviews Neuroscience 12, 217–230 (2011). 10.1038/nrn3008

40 Mishkin, M. & Ungerleider, L. G. Contribution of striate inputs to the visuospatial functions of parieto-preoccipital cortex in monkeys. Behav Brain Res 6, 57–77 (1982). 10.1016/0166-4328(82)90081-x

41 Mishkin, M., Ungerleider, L. G. & Macko, K. A. Object vision and spatial vision: two cortical pathways. Trends in Neurosciences 6, 414–417 (1983). 10.1016/0166-2236(83)90190-X

42 Hebart, M. N. & Hesselmann, G. What Visual Information Is Processed in the Human Dorsal Stream? The Journal of Neuroscience 32, 8107–8109 (2012). 10.1523/jneurosci.1462-12.2012

43 Norman, J. Two visual systems and two theories of perception: An attempt to reconcile the constructivist and ecological approaches. Behavioral and Brain Sciences 25, 73–96 (2002). 10.1017/S0140525X0200002X

44 Kozlovskiy, S. R., Anton (2021). “How Areas of Ventral Visual Stream Interact When We Memorize Color and Shape Information”. Advances in Intelligent Systems and Computing. Springer-Nature. 1358 (95–100).

45 Schneider, G. E. Two Visual Systems. Science 163, 895–902 (1969). doi:10.1126/science.163.3870.895

46 Gallivan, J. P., Chapman, C. S., Wolpert, D. M. & Flanagan, J. R. Decision-making in sensorimotor control. Nature Reviews Neuroscience 19, 519–534 (2018). 10.1038/s41583-018-0045-9

47 Chang, E. H. et al. The role of myelination in measures of white matter integrity: Combination of diffusion tensor imaging and two-photon microscopy of CLARITY intact brains. Neuroimage 147, 253–261 (2017). 10.1016/j.neuroimage.2016.11.068

48 Kochunov, P. et al. Fractional anisotropy of water diffusion in cerebral white matter across the lifespan. Neurobiology of Aging 33, 9–20 (2012). 10.1016/j.neurobiolaging.2010.01.014

49 Bukkieva, T. et al. Microstructural Properties of Brain White Matter Tracts in Breast Cancer Survivors: A Diffusion Tensor Imaging Study. Pathophysiology 29, 595–609 (2022). 10.3390/pathophysiology29040046

50 Murray, L. et al. Sex differences in functional network dynamics observed using coactivation pattern analysis. Cognitive Neuroscience 12, 120–130 (2021). 10.1080/17588928.2021.1880383

51 Ferreira, L. K. et al. Aging Effects on Whole-Brain Functional Connectivity in Adults Free of Cognitive and Psychiatric Disorders. Cerebral Cortex 26, 3851–3865 (2016). 10.1093/cercor/bhv190

52 Griffa, A. et al. Brain connectivity alterations in early psychosis: from clinical to neuroimaging staging. Translational Psychiatry 9, 62 (2019). 10.1038/s41398-019-0392-y

53 Maximo, J. O., Cadena, E. J. & Kana, R. K. The implications of brain connectivity in the neuropsychology of autism. Neuropsychol Rev 24, 16–31 (2014). 10.1007/s11065-014-9250-0

54 Wang, M. et al. Disrupted functional brain connectivity networks in children with attention-deficit/hyperactivity disorder: evidence from resting-state functional near-infrared spectroscopy. Neurophotonics 7, 015012 (2020). 10.1117/1.NPh.7.1.015012

55 Yeh, F. C., Verstynen, T. D., Wang, Y., Fernandez-Miranda, J. C. & Tseng, W. Y. Deterministic diffusion fiber tracking improved by quantitative anisotropy. PLoS One 8, e80713 (2013). 10.1371/journal.pone.0080713

56 Kral, T. R. A. et al. Neural correlates of video game empathy training in adolescents: a randomized trial. npj Science of Learning 3, 13 (2018). 10.1038/s41539-018-0029-6

57 Kühn, S., Gleich, T., Lorenz, R. C., Lindenberger, U. & Gallinat, J. Playing Super Mario induces structural brain plasticity: gray matter changes resulting from training with a commercial video game. Molecular Psychiatry 19, 265–271 (2014). 10.1038/mp.2013.120

58 Pasqualotto, A., Parong, J., Green, C. S. & Bavelier, D. Video Game Design for Learning to Learn. International Journal of Human–Computer Interaction 39, 2211–2228 (2023). 10.1080/10447318.2022.2110684

59 Cunningham, E. G. & Green, C. S. (Oxford University Press, 2023).

60 van Polanen, V. & Davare, M. Interactions between dorsal and ventral streams for controlling skilled grasp. Neuropsychologia 79, 186–191 (2015). 10.1016/j.neuropsychologia.2015.07.010

61 Kersey, A. J., Clark, T. S., Lussier, C. A., Mahon, B. Z. & Cantlon, J. F. Development of Tool Representations in the Dorsal and Ventral Visual Object Processing Pathways. Cereb Cortex 26, 3135–3145 (2016). 10.1093/cercor/bhv140

62 Puderbaugh, M. & Emmady, P. D. in StatPearls (StatPearls Publishing Copyright © 2024, StatPearls Publishing LLC., 2024).

63 Wong, W.-w., et al. Effects of visual attention modulation on dynamic effective connectivity and visual fixation during own-face viewing in body dysmorphic disorder. medRxiv, 2021.2002.2015.21249769 (2021). 10.1101/2021.02.15.21249769

64 Xia M, W. J., He Y (2013) BrainNet Viewer: A Network Visualization Tool for Human Brain Connectomics. PLoS ONE 8(7): e68910. 10.1371/journal.pone.0068910.

65 Giacalone, M., Agata, Z., Cozzucoli, P. C. & Alibrandi, A. Bonferroni-Holm and permutation tests to compare health data: methodological and applicative issues. BMC Medical Research Methodology 18, 81 (2018). 10.1186/s12874-018-0540-8

66 Holm, S. A simple sequentially rejective multiple test procedure. Scandinavian journal of statistics, 65–70 (1979).

67 Bastos, A. M. & Schoffelen, J.-M. A Tutorial Review of Functional Connectivity Analysis Methods and Their Interpretational Pitfalls. Frontiers in Systems Neuroscience 9 (2016). 10.3389/fnsys.2015.00175

68 Friston, K. J. Functional and effective connectivity in neuroimaging: A synthesis. Human Brain Mapping 2, 56–78 (1994). 10.1002/hbm.460020107

69 Rosch, K. S. & Mostofsky, S. in Handbook of Clinical Neurology Vol. 163 (eds Mark D’Esposito & Jordan H. Grafman) 351-367 (Elsevier, 2019).

70 Eickhoff, S. B. & Müller, V. I. in Brain Mapping (ed Arthur W. Toga) 187-201 (Academic Press, 2015).

71 Dhamala, M., Liang, H., Bressler, S. L. & Ding, M. Granger-Geweke causality: Estimation and interpretation. Neuroimage 175, 460–463 (2018). 10.1016/j.neuroimage.2018.04.043

72 Dhamala, M., Rangarajan, G. & Ding, M. Estimating Granger Causality from Fourier and Wavelet Transforms of Time Series Data. Physical Review Letters 100, 018701 (2008). 10.1103/PhysRevLett.100.018701

73 Dhamala, M., Rangarajan, G. & Ding, M. Analyzing information flow in brain networks with nonparametric Granger causality. Neuroimage 41, 354–362 (2008). 10.1016/j.neuroimage.2008.02.020

74 Yeh FC, P. S. F. D. M. A., Yoshino M Fernandez-Miranda JC Vettel JM, Verstynen T. Population-averaged atlas of the macroscale human structural connectome and its network topology. Neuroimage (2018). doi: 10.1016/j.neuroimage.2018.05.027.

75 Yeh, F. C., Wedeen, V. J. & Tseng, W. Y. Generalized q-sampling imaging. IEEE Trans Med Imaging 29, 1626–1635 (2010). 10.1109/tmi.2010.2045126

76 Yeh, F. C. & Tseng, W. Y. NTU-90: a high angular resolution brain atlas constructed by q-space diffeomorphic reconstruction. Neuroimage 58, 91–99 (2011). 10.1016/j.neuroimage.2011.06.021

77 Babaeeghazvini, P., Rueda-Delgado, L. M., Gooijers, J., Swinnen, S. P. & Daffertshofer, A. Brain Structural and Functional Connectivity: A Review of Combined Works of Diffusion Magnetic Resonance Imaging and Electro-Encephalography. Frontiers in Human Neuroscience 15 (2021). 10.3389/fnhum.2021.721206

78 Sporns, O. Structure and function of complex brain networks. Dialogues Clin Neurosci 15, 247–262 (2013). 10.31887/DCNS.2013.15.3/osporns

79 Soares, J., Marques, P., Alves, V. & Sousa, N. A hitchhiker’s guide to diffusion tensor imaging. Frontiers in Neuroscience 7 (2013). 10.3389/fnins.2013.00031

80 Sammer, G., Neumann, E., Blecker, C. & Pedraz-Petrozzi, B. Fractional anisotropy and peripheral cytokine concentrations in outpatients with depressive episode: a diffusion tensor imaging observational study. Scientific Reports 12, 17450 (2022). 10.1038/s41598-022-22437-0

81 Ahn, S. & Lee, S.-K. Diffusion Tensor Imaging: Exploring the Motor Networks and Clinical Applications. Korean journal of radiology : official journal of the Korean Radiological Society 12, 651–661 (2011). 10.3348/kjr.2011.12.6.651

